# Time- and space-resolved regulatory circuits in *Arabidopsis thaliana*

**DOI:** 10.1101/2025.10.23.684094

**Authors:** Sanja Zenker, Katharina Schiller, Andrea Bräutigam

## Abstract

General metabolism and responses to internal or external signals are tightly regulated in plants. We hypothesize that a network of intermediate regulatory proteins including transcription factors, kinases, and E3 ligases connect input signals like light, temperature, circadian clock, and ontogenetic pathways to output pathways. We therefore investigate the transcriptional dynamics of these regulators.
RNA sequencing was performed on three-week-old *Arabidopsis thaliana* Col-0 rosettes grown under 12h/12h light/dark conditions sampled every two hours over 24 hours to match publicly available single cell data of same age plants. Both were analyzed to identify the abundance of intermediate regulators in time and space.
Intermediate regulators are of significantly lower abundance compared to other transcripts in the Arabidopsis transcriptome. More than half of expressed kinases, E3 ligases and transcriptions factors vary in either time, space or both in mature leaves.
Dynamic expression patterns of regulators allow plants to maintain tightly regulated metabolism while providing sufficient room for specific stress responses. High plasticity of the Arabidopsis transcriptome highlights the importance of considering sampling time-of-day and cellular resolution for experiments.

**One sentence summary:** Expression patterns in time and space of kinases, E3 ligases and transcription factors in *Arabidopsis thaliana* provide ample room for dynamic and specific regulation of metabolism and stress responses.

## Introduction

*Arabidopsis thaliana* contains a large proportion of genes with regulatory functions in its genome. Regulatory proteins include transcription factors (TFs), which occupy 6-7% of total genes (Riechmann *et al*. 2000; Petroll *et al*. 2025), kinases, which account for 4% (Zulawski *et al*. 2014), and the E3 ligase family which has more than 1,400 members (Vierstra 2009). Closely related TFs are often functionally redundant to each other (Leivar, Monte, Oka, *et al*. 2008) by sharing binding motifs and binding sites (Zenker *et al*. 2025). Kinases phosphorylate targets comparatively unspecifically *in vitro* (Baena-González *et al*. 2007) and can act redundantly (Tena *et al*. 2011). Expression changes in space and time have been suggested to account for specificity in the face of apparent redundancy.

Mature Arabidopsis leaves contain multiple cell types, guard cells, stomata and pavement cells in the epidermis, spongy and palisade mesophyll cells, and bundle sheath cells surrounding phloem and xylem in the veins (Lee *et al*. 2025). The cells in these tissues share features, often called housekeeping, such as DNA maintenance, protein synthesis, and others. But they also have specific features such as a cuticula maintained by epidermal cells or transport capacity needed in the phloem (Lucas *et al*. 2013; Lewandowska *et al*. 2020). Combinatorial effects of regulatory proteins are hypothesized to determine cell specificity. The differences in regulatory proteins also lead to different output to stresses in different cell types (Wang *et al*. 2025; Tenorio Berrío *et al*. 2025). In addition to this spatial diversity, cells in mature Arabidopsis leaves are governed by diurnal changes cued by light, temperature, and the circadian clock. Global transcriptomic analyses based on microarrays identify ∼6-50% of expressed genes as diurnal (Harmer *et al*. 2000; Bläsing *et al*. 2005) and recent RNA-seq based analyses 61% (Redmond *et al*. 2025). The significance of the diurnal changes in transcripts is unclear since proteomics experiments have determined that diurnal changes in the Arabidopsis leaf transcriptome and proteome do not match as most quantifiable proteins do not oscillate (Baerenfaller *et al*. 2012). The median protein half-life is 6.3 days for quantifiable proteins (Li *et al*. 2017). The restriction “quantifiable” is important in these assessments since only proteins with sufficient abundance can be reliably quantified in either approach. Proteins with regulatory functions are frequently underrepresented in these analyses (Baerenfaller *et al*. 2012; Li *et al*. 2017). We hypothesized that genes with regulatory functions are expressed at lower levels leading to reduced transcript per million contributions to the transcriptome.

Metabolism in general and responses to stresses require the regulation of proteins in time and space. Low protein degradation rates make it unlikely that protein turnover of enzymes in diurnal patterns governs the abundance of enzymes and therefore the flux (Schiller *et al*. 2025). In contrast, modulation of enzyme activity via kinases may tune activity throughout the day. For example, the enzyme phospho*enol*pyruvate carboxylase (PEPC) which produces oxaloacetate from PEP is regulated by PEPC kinase which is tightly regulated in diurnal patterns to tune day and nighttime metabolism to different photosynthetic subtypes such as C3, C4, and CAM photosynthesis (Hartwell *et al*. 1999; Aldous *et al*. 2014). For many response pathways, the ability to respond is different in time. Responses to internal (Lee *et al*. 2016), abiotic (Allen *et al*. 2006; Li *et al*. 2019) and biotic signals (Bhardwaj *et al*. 2011) are gated by the circadian clock. Protein synthesis is imbalanced between day and night (Piques *et al*. 2009; Duncan & Millar 2022) and, since protein levels are fairly stable (Li *et al*. 2017), protein degradation is likely highly regulated (Vierstra 2009).

Cell identity is established during ontogeny and is, with the exception of regenerative processes, stable. For example, once mesophyll is differentiated, veins can no longer be formed (Scarpella *et al*. 2004). Prior to mesophyll differentiation, vein formation is initiated by a series of transcriptional events (Scarpella *et al*. 2004). We hypothesize that after these initial determinants during ontogeny, intermediate regulatory proteins maintain the necessary processes in mature leaves. Similarly, diurnal expression is controlled by top-level regulation systems. The core circadian clock in Arabidopsis consists of a feedback circuit between differently phased regulators ensuring 24-h oscillation by transcriptional control (Staiger *et al*. 2013). Components of the core clock are also regulated post-transcriptionally, for example Circadian Clock Associated 1 (CCA1) is phosphorylated by Casein Kinase 2 (CK2) (Daniel *et al*. 2004) and Timing of CAB Expression 1 (TOC1) can be ubiquitinylated by F-box protein Zeitlupe (ZTL) (Harmon *et al*. 2008). Light and temperature also cue diurnal expression. Light is perceived through red and blue light receptors and its signal is transmitted to transcriptional regulators such as Phytochrome Interacting Factor (PIF) proteins and Elongated Hypocotyl 5 (HY5) for red light (Leivar, Monte, Al-Sady, *et al*. 2008; Van Gelderen *et al*. 2018) and to Target of EAT1 (TOE1) for blue light receptors (Du *et al*. 2020). Temperature also feeds into PIF TFs, e.g. PIF7 (Chung *et al*. 2020). Evidence for several protein families suggests that differential expression patterns in time and space are common. Many TFs downstream of the core clock show diurnal expression patterns, for example members of the C2-like BBX-type family often peak shortly after dawn (Balcerowicz *et al*. 2021). Downstream TFs have also been identified as locally expressed, for example transcripts of multiple members of the NAC and R2R3 MYB family have been shown to localize in the phloem (Zhao *et al*. 2005) or the TF FAMA in guard cells (Ohashi-Ito & Bergmann 2006).

We hypothesize that output pathways such as general metabolism and responses to signals are connected to a network of intermediate regulatory proteins including TFs, kinases, and E3 ligases, which in turn receive signals from initial top-level regulators downstream of sensors. To contrast and compare expression patterns in space and in time, we generated a 24 hour diurnal leaf transcriptomic dataset sampled every two hours that is developmentally matched to the single cell leaf transcriptome of 21-day old Arabidopsis rosettes (Lee *et al*. 2025). The data shows that, of the 770 expressed kinases, 509 vary in time, space, or both in a mature leaf. The majority peak at zt13 and 32.74% of rhythmic kinases peak between zt13 and zt15. The E3 ligases for protein degradation also vary to a large degree with 420 of 732 expressed genes varying in time, space or both. Similarly, of 1,121 expressed TFs, 708 vary in time and/or space. We conclude that at a plant age of 21 days in mature aerial tissues, a majority of regulatory transcripts vary to large degrees enabling the time- and space-specific regulation of metabolism and stress response. The regulatory space not only includes transcriptional regulation, which is evident in the large proportion of the transcriptomic variation of TFs and target genes, but also post-transcriptional regulation as evidenced by dynamically expressed low-abundance kinases and E3 ligases.

## Methods

### Growth conditions and sampling

*Arabidopsis thaliana* Col-0 seeds were surface sterilized using Ethanol and stratified for 2 days at 4°C in the dark. Seeds were transferred on ½ MS Agar plates (0.8%) without addition of any sugar and grown for three weeks in a climate chamber with a 12 h/12 h day/night cycle: 22°C, 120 µE and 18°C 0 µE. Relative humidity was constant at 50%. Above ground material from three whole plates was harvested every 2 hours as three biological replicates by flooding the plate with liquid nitrogen. Sampling was started at Zeitgeber time 2 (zt2) and ended with a repetition of zt2* the next day. Zt12 was sampled shortly before darkness and zt0 directly before the lights turned on. Transitions were immediate with no light gradient applied. RNA was extracted using Qiagen RNeasy Plant Mini Kit and a 15 minute on-column DnaseI digestion. RNA library was prepared using Illumina XLEAP chemistry and sequenced as single strands on a NextSeq2000 yielding 6.5 to 12.7 million reads per sample.

### Data analysis

Raw reads were aligned to the TAIR10 Arabidopsis transcriptome of only primary transcripts with kallisto v0.44.0 (Bray *et al*. 2016) with the parameters “--single −l 200 -s 20” resulting in mapping rates of 92.4% to 94.8%. Further downstream analyses were performed using R v4.5.1 (R Core Team 2024) and the tidyverse (Wickham *et al*. 2019). A PCA was computed with prcomp on scaled and centered expression data. To determine rhythmic transcripts, JTK_CYCLE (Hughes *et al*. 2010) was used on expressed transcripts, here defined as >1 tpm in all three replicates of at least one time point. A target period length of 24 hours was set and transcripts with Benjamini-Hochberg adjusted p-value <0.01 and an amplitude >1 were considered diurnal. GO term enrichments were performed using topGO (Adrian Alexa 2017) on all genes peaking at a given Zeitgeber time. To analyze spatial resolution, data from Lee *et al*. (2025) was reanalyzed. The Seurat object of single cell RNA-seq data of 21d old rosettes was accessed from GSE226097 and cell types were assigned based on Lee *et al*. (2025) Supplemental Table 2. Cell type specific expressed genes were identified using FindAllMarkers from the Seurat v5.3.0 R package (Hao *et al*. 2024) with min.pct=0.1, logfc.threshold=0.25 and onl.pos=T. Genes with a p-value<0.05 in not more than two cell types were defined as spatially specific. For the fraction of cells with expression, a gene was considered expressed if at least one transcript was detected in a cell.

To annotate the intermediate regulators, TFs in *A. thaliana* were retrieved from the TAPscan v4 database (Petroll *et al*. 2025). Proteins with kinase domains were selected via HMMER search using the HMM models for Pfam domains Pkinase (PF00069) and PK_Tyr_Ser-Thr (PF07714) and default settings (Finn *et al*. 2011). Subclade annotations of kinases were retrieved from Zulawski *et al*. (2014). E3 ubiquitin ligases mediating substrate specificity, namely HECT, RING, F-box, U-box, BTB, DDB1 proteins and APC complex recognition subunits (Vierstra 2009) were retrieved from Mazzucotelli *et al*. (2006). Joined heatmaps of diurnal and spatial data were generated with the R package ComplexHeatmap (Gu 2022) from z-scores of diurnal genes with Euclidean distance and Ward’s Clustering. All processed data is available in Table S1.

## Results

To test the diurnal dataset for overall expression patterns, a principal component analysis (PCA) and phasing of transcripts were analyzed. The PCA partitioned 38.91% of the variation in the first two components (Figure 1A). Transcriptomic changes are stable across biological replicates and follow a 24h-period reaching zt2 again at zt2* the following day. The pattern resembles a circle in the first two principal components (Figure 1A). Samples taken in the dark are clearly separated from those in the light (Figure 1A). Out of 27,206 nuclear protein coding genes, 18,337 (67.4%) genes are detectable at >1 tpm at least at one timepoint in aerial parts of 21-day old Arabidopsis plants, of which 62% (11,364) are detected as diurnal (p_adj_ <0.01; amplitude >1) within a 24-hour period by JTK_CYCLE (Figure 1B, Table S2). Regulatory transcripts are of significantly lower abundance compared to all transcripts (p=1.01e-26 for TFs, p=1.34e-15 for kinases, and p=2.53e-32 for E3 ligases, Figure 1C). The largest number of diurnal transcripts peak at the beginning and middle of the night (Figure 1D). Consequently, genes with peak expression at zt15, zt16, and zt17 enrich in GO terms related to mRNA metabolic processes (Figure 1D, Table S3). The GO term photosynthesis is enriched at zt1-zt4 while response to light stimulus is already enriched one hour before lights turned on at zt23 (Figure 1D).

**Figure 1.**
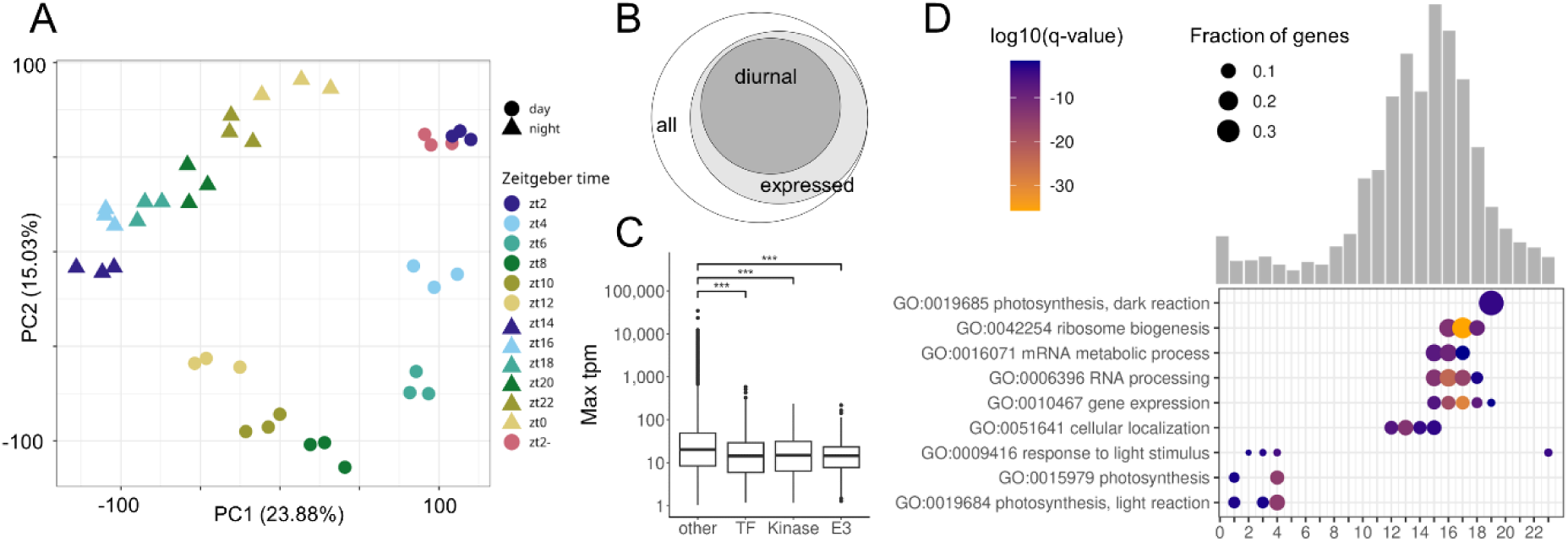
Global diurnal expression patterns in the Arabidopsis transcriptome. **A**: PCA of 13 sampled timepoints with three biological replicates each. Colored by Zeitgeber time with shape indicating day and night. **B**: Euler diagram showing the proportions of expressed and diurnal transcripts in Arabidopsis leaves. **C**: Boxplots of maximal expression levels per gene per day. Wilcoxon test compares kinases, E3 ligases and TFs to all other genes (***=p<0.001). **D**: Number of diurnal transcripts grouped by their peak phase as defined by JTK_CYCLE (Table S2) indicated by bars. Selected enriched GO terms (q-value<0.05) per phase are shown with size indicating fraction of phased genes in GO term and color encoding log10 q-value.

### Kinases in time and space

To test kinases for their variation in time and space (Figure 2), they were grouped into 559 receptor kinases and 381 soluble kinases based on Zulawski *et al*. (2014). We find 770 of 1,051 kinases expressed and 452 of those (58.7%) to be diurnal (Figure 2B). Subsets of 209 receptor kinases (37.39%) and 205 soluble kinases (53.8%) are diurnal (p_adj_ <0.01; amplitude >1). The heatmap of diurnal transcript levels (Figure 2C) shows that different kinases peak in abundance during different times of the day, but the majority of 35.84% (162) peak around the beginning of the night (zt13-zt15). This is true for both subclades of kinases, receptor kinases and soluble kinases (Figure 2C). Only 8.8% of the diurnal soluble kinases peak in the first eight hours of the day but 21.95% peak in the final four hours of the day. Of the diurnal receptor kinases, 3.45% peak in the first four hours of the day, 8.13% peak around noon, and 25.36% peak in the final four hours of the day. Soluble kinases and receptor kinases which peak at the beginning of the night frequently remain elevated in abundance throughout the night compared to their daytime abundance (Figure 2C). A small group of kinases peak during the end of the night and continue into the day (Figure 2C). Among the soluble kinases, ABA-activated kinase Open Stomata 1 (OST1; AT4G33950) which phosphorylates Slow Anion Channel-Associated 1 (SLAC1) peaks at zt17 and is expressed ubiquitously in all cell types (Figure S1). Mitogen-activated Protein Kinase 3 (MPK3; AT3G45640) and MPK6 (AT2G43790) involved in pathogen-associated molecular pattern recognition peak at zt10 and zt15 respectively (Figure S1). MPK3 is significantly expressed in trichome cells but MPK6 is not cell type specific (Figure S1). PEPC kinase (AT1G08650) peaks at zt7 and is not cell type specific (Figure S1). The receptor kinase Brassinosteroid Insensitive 1 (BRI1; AT4G39400) involved in brassinosteroid signaling peaks at zt13 and is cell type specific in the phloem and trichome (Figure S1). Receptor kinase Flagellin-sensitive 2 (FLS2; AT5G46330) peaks at zt13 and is significantly differentially expressed in the phloem (36% of cells), but also found in 19-26% of cells of other types (Figure S1). The phloem overall has the highest amount of spatially specific genes with 2,701 genes, followed by 563 in the trichome (Table S4). Similar to these single gene examples, the majority of leaf kinases are dynamic in time, space or in both. If spatial specificity is defined as differentially expressed in a maximum two cell types, 205 expressed kinases are spatially specific (p_adj_ <0.01; Table S4). A subset of 148 of the 452 diurnal kinases are diurnal and spatial (Figure 2B). In summary, 66.1% of the kinases expressed in 21-day old aerial parts of Arabidopsis are dynamic in time, space or both. Remarkably, on/off behaviour was detected in 74 of the 770 expressed kinases with an average expression of <1 tpm at at least one time point during the diurnal time-course.

**Figure 2.**
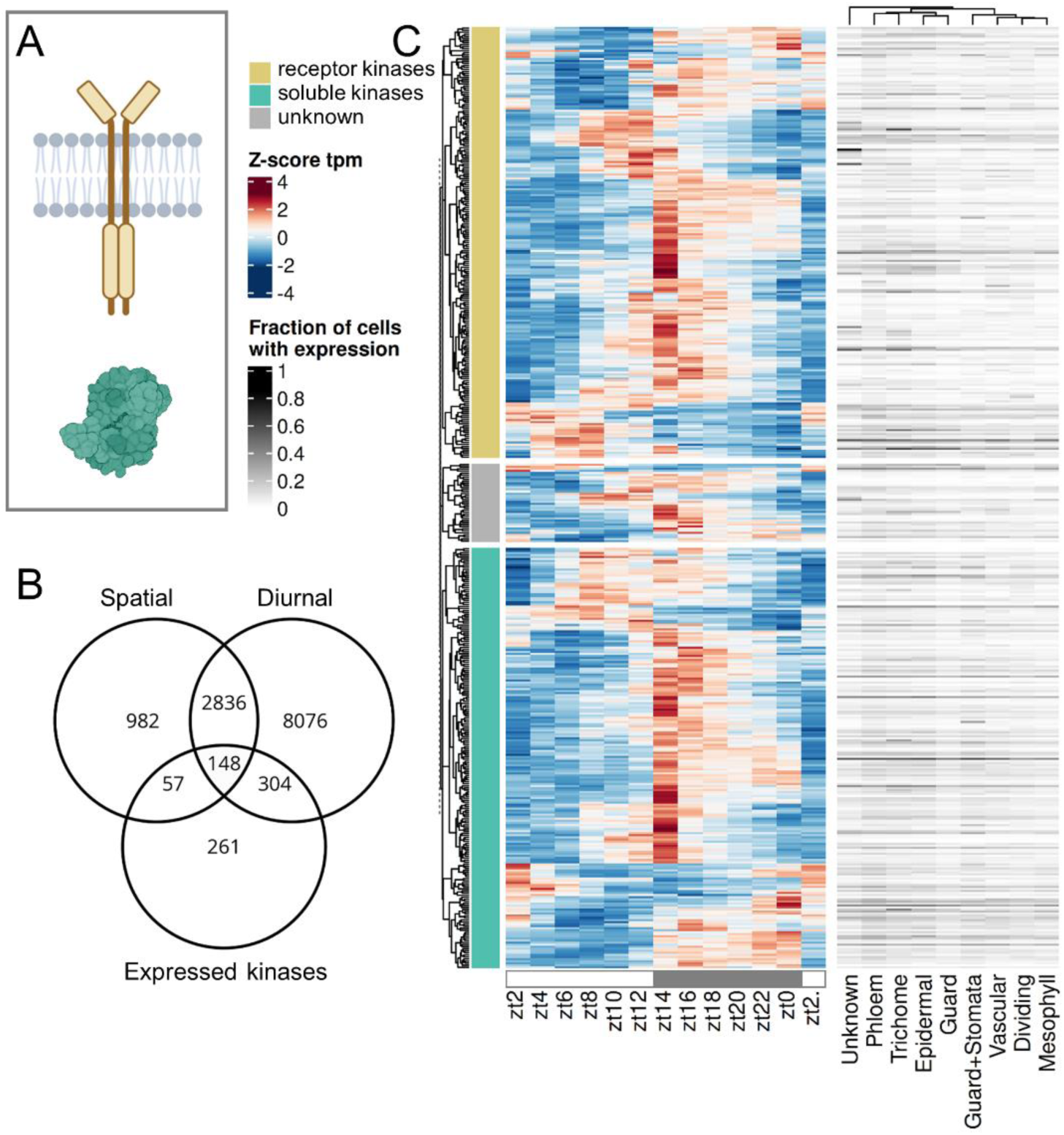
Expression patterns of kinases. **A**: Schematic visualization of a membrane located and a soluble kinase as the two main subtypes (Zulawski *et al*. 2014). **B**: Venn diagram showing the overlap between diurnal, spatial cell type specific genes and expressed kinases. **C**: Heatmap showing expression (z-score tpm) of all diurnal kinase transcripts and fraction of cells with expression for each cell type (Lee *et al*. 2025) grouped by subclades (Zulawski *et al*. 2014).

### E3 ligases in time and space

Single protein E3 ligases (e.g. RING; Figure 3A) and the target specificity mediating proteins from E3 ligase complexes (hereafter E3 ligases) were tested for their variation in time and space (Figure 3). Seven HECT, 486 RING, 49 U-box, 81 BTB, 5 DDB1, 700 F-box and 10 other nuclear protein coding genes were annotated based on Mazzucotelli *et al*. (2006). In total, we find 732 of 1,338 E3 ligases expressed in 21-day old rosettes and 380 (51.91%) to be diurnal (p_adj_ <0.01; amplitude >1, Figure 3B). Different E3 ligases peak in transcript abundance during different times of the day (Figure 3C), but the majority of 28.35% peak around the beginning of the night (zt13-zt15). A small subset of 3.41% of the diurnal E3 ligases peak in the first four hours of the day, 4.46% peak around noon, and 21.8% peak in the final four hours of the day. Transcript levels of F-box, RING and BTB proteins which peak at the beginning of the night frequently remain elevated in abundance throughout the night, which is less observed for U-box proteins (Figure 3C). Specifically, within the largest subgroup of F-box proteins, 2.67% of the diurnal transcripts peak in the first four hours of the day, as well as at noon, and 18.75% peak in the final four hours of the day. The auxin receptor F-box protein Transport Inhibitor Response 1 (TIR1; AT3G62980) peaks at zt17 and is cell type specific in phloem and epidermal cells in 21-day old rosettes (Figure S1). The jasmonate receptor Coronatine Insensitive 1 (COI1; AT2G39940) is not diurnal but is cell type specific in the phloem (Figure S1). The E3 ligase Constitutive Photomorphogenic 1 (COP1; AT2G32950) is also not diurnal but cell type specific in the phloem and trichome (Figure S1). Transcripts of High Expression of Osmotically Responsive Genes 1 (HOS1; AT2G39810), which is a negative regulator of cold tolerance, peak at zt16 and are detected in ∼60% of guard and stomatal cells and significantly enrich for this cell type (Figure S1, Table S1). We find 17.35% of transcripts of E3 ligase genes spatially specific and a subset of 87 of the 732 diurnal transcripts are not only diurnal but also spatially specific (Figure 3B). In summary, 57.38% of the transcripts for proteins mediating the specificity in protein degradation expressed in 21-day old aerial parts of Arabidopsis are dynamic in time or space or both. An on/off behavior was detected in 66 of the 732 expressed E3 ligases.

**Figure 3.**
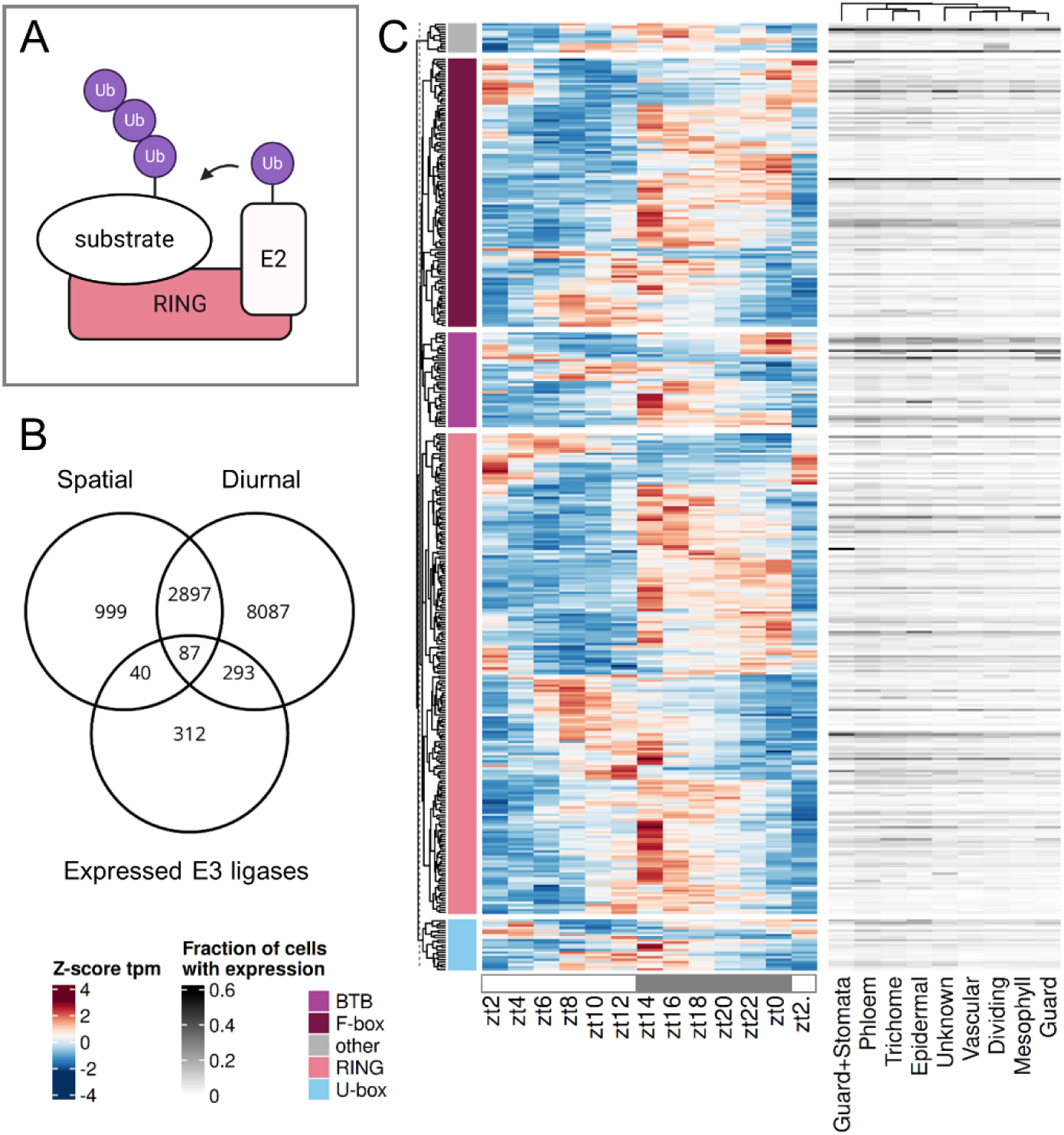
Expression patterns of E3 ligases. **A**: Schematic visualization of the substrate specific ubiquitinylation by a RING-type ligase. **B**: Venn diagram showing the overlap between diurnal, spatial cell type specific genes and expressed E3 ligases. **C**: Heatmap of all diurnal transcripts (z-score tpm) over time and fraction of cells with expression for each cell type (Lee *et al*. 2025). Annotations of the four largest subgroups are color coded and proteins of smaller subgroups are combined under other.

### Transcription factors in time and space

Transcription factors (TFs) were tested for their variation in time and space (Figure 4). Only DNA-binding TFs were considered (Figure 4A). We find 1,121 of 1,725 DNA-binding TFs expressed in aerial tissues and 649 (57.89%) to be diurnal (Figure 4B). The heatmap shows that transcript abundance of different TFs peak during different times of the day (Figure 3C). 6.47% of the diurnal TFs peak in the first four hours of the day, 6.16% peak around noon, and 20.19% peak in the final four hours of the day while 32.67% peak in the first four hours of the night, 23.42% at midnight, and 11.09% at the end of the night (Figure 4C). A small group of TFs peak towards the end of the night and into the day (Figure 4C). The top-level circadian clock regulators peak at zt22 (LHY), zt23 (CCA1), and at zt12 (TOC1) and are generally expressed in all cell types (Figure S2, Table S1), but LHY and CCA1 are also differentially expressed in the phloem (Figure S1).

**Figure 4.**
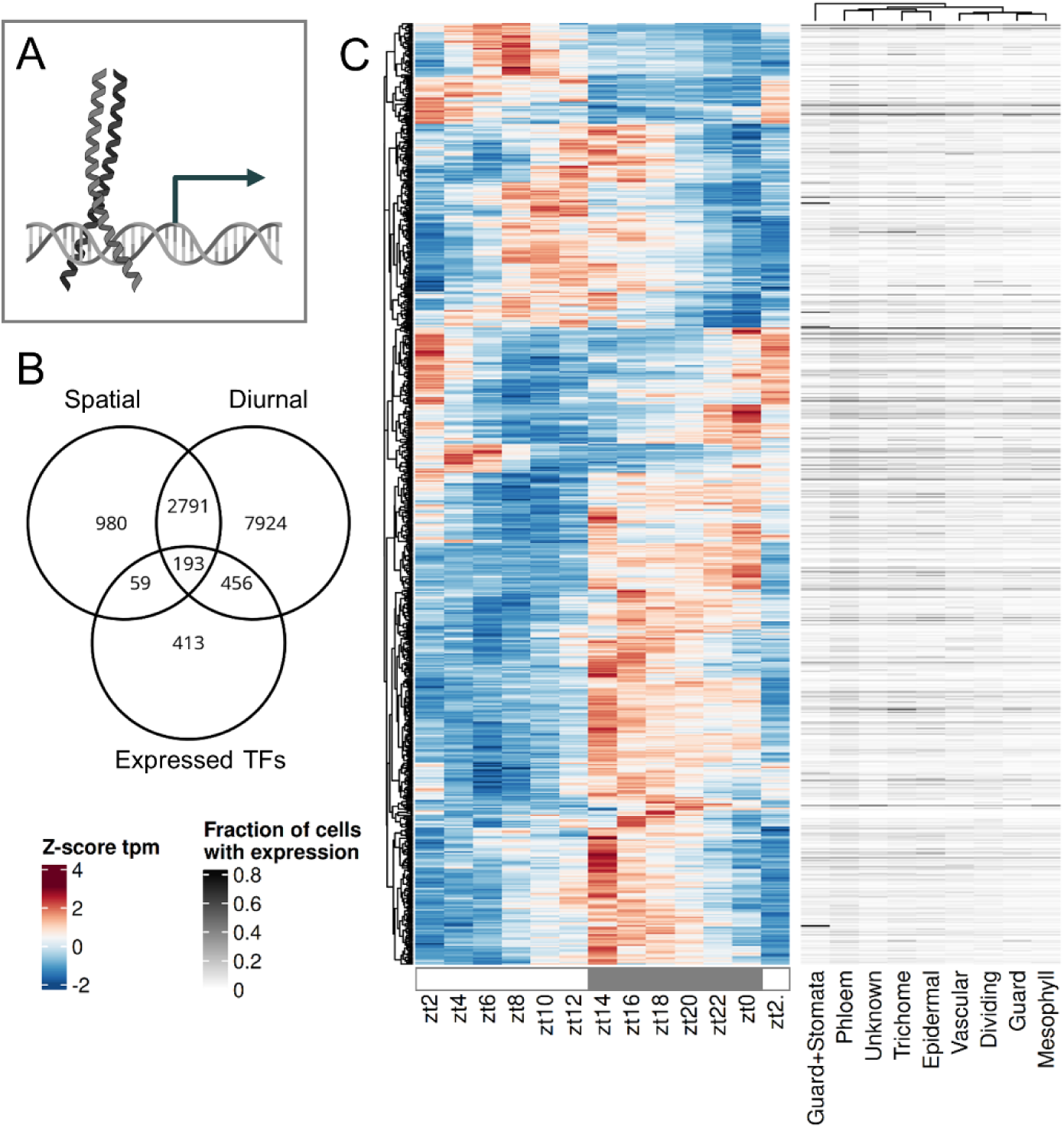
Expression patterns of transcription factors. **A**: Schematic visualization of a DNA-binding TF, in this case a bZIP-type. **B**: Venn diagram showing the overlap between diurnal, spatial cell type specific transcripts and expressed TFs. **C**: Heatmap of expression (z-score tpm) of all diurnal transcripts over time and fraction of cells with expression for each cell type (Lee *et al*. 2025).

The TF FAMA (AT3G24140) is a known regulator of guard cell fate which peaks at zt14 and is generally more expressed during the night (Figure S1). Transcripts significantly enrich in the guard and stomata lineage cluster, where at least one transcript is detected in 83% of cells (Figure S1, Table S1). The photosynthetic transcription factor Golden2-like 1 (GLK1; AT2G20570) peaks at zt6 and is enriched in the phloem (Figure S1). Similar to these TFs, the majority of transcripts for TFs are dynamic in time, in space or in both. If spatially specific is defined as in maximally two cell types, 22.48% of TFs are spatial (Figure 4B). A subset of 193 TF transcripts are both diurnal and spatial (Figure 4B). In summary, 63.16% of the transcripts for TFs expressed in 21-day old aerial parts of Arabidopsis are dynamic in time or space or both. Transcripts for TFs show on/off behavior with 187 of the 1,121 expressed ones with an average expression of <1tpm at at least one time point during the diurnal time-course.

## Discussion

The data for diurnal expression patterns of the Arabidopsis rosettes over 24 hours grown under 12h light/12h dark growth conditions (Table S1) combined with the matched single cell atlas (Lee *et al*. 2025) enables the analysis of the intermediate regulatory layer (Figures 2, 3, 4). The analysis detects 62% of transcripts as diurnal (Figure 1B, Table S2) which is comparable to Redmond *et al*. (2025) and broadly comparable to microarray datasets (Harmer *et al*. 2000; Bläsing *et al*. 2005). If cut-off values for diurnal rhythmicity are relaxed (see methods) an even larger proportion of the transcriptome would be considered diurnal. This proportion massively exceeds the known direct targets of the circadian clock (Nagel *et al*. 2015; Ezer *et al*. 2017). This observation is likely explained by the large proportion of diurnal TFs (Figure 4). Given that many TFs share target binding motifs and sites (Zenker *et al*. 2025) a large proportion of TFs with diurnal behavior will result in a large proportion of the total transcriptome with some degree of rhythmic expression (Figure 1B).

PEPC kinase is one example for a regulatory kinase which directs metabolic flow (Schiller & Bräutigam 2021). The kinase has on/off behavior in CAM plants being on during the night (Hartwell *et al*. 1999). In stark contrast, transcripts peak during the day in the C3 plant Arabidopsis at zt7 but are not on/off (Figure S1, Table S1), which is characteristic for a C3 plant (Fontaine *et al*. 2002) and highlights the different needs of regulation for PEPC in a C3 background. Seventy-four kinases show on/off behavior in Arabidopsis aerial tissue during a day/night time-course similar to the PEPC kinase in CAM plants and may perform similar roles for different processes and fluxes. FLS2, the pathogen associated pattern recognition sensor which recognized flagellin, surprisingly peaks during the early night but does not show on/off behavior (Figure S1, Table S1). Given that stomata close during the night, this expression likely does not relate to function. This behavior highlights that a proportion of diurnal changes may not be related to function of the protein itself but is a function of the large proportion of transcription factors with diurnal behavior (Figure 4) and their partial promiscuity (Zenker *et al*. 2025). OST1 extended roles beyond stomatal regulation (Mustilli *et al*. 2002) is highlighted by its expression throughout the leaf cell types (Figure S1, Table S1). This suggests that some but not all observed patterns for kinase transcript abundances (Figure 2) have functional significance.

Timed experiments with protein synthesis have suggested that protein degradation is regulated diurnally (Piques *et al*. 2009; Ishihara *et al*. 2015). Indeed, there is very large potential for targeted degradation as 1,338 target specificity mediating proteins from E3 ligase complexes are annotated (Mazzucotelli *et al*. 2006; Vierstra 2009). Of these, 380 have diurnal behavior but the majority of them peaks at the beginning of the night (Figure 3C). To balance protein synthesis as suggested (Ishihara *et al*. 2015), a peak of targeted degradation during the day is required. The balancing function is thus carried out by a minority of E3 ligases (Figure 3C). The majority peak of E3 ligases at the beginning of the night coincides with the majority peak of kinases at the beginning of the night (Figures 2C, 3C) which may suggest that the kinase based signaling may be subject to rapid and targeted turnover by E3 ligases. The fact that all signaling proteins tested have significantly lower transcript abundance (Figure 1C) combined with the observation that protein abundance positively correlates with transcript abundance (Mergner *et al*. 2020) makes signaling proteins subject to a potentially rapid rate of turnover undetected by proteomics experiments as these quantify mostly abundant proteins.

Using this time and space resolved data, it is possible to confirm known spatiotemporal expression patterns but also define a distinct pattern or absence thereof for other genes. For example, FAMA is a known determinant of guard cell fate and generally used to annotate the guard and stomata lineage clusters in single cell analyses (Ohashi-Ito & Bergmann 2006; Lee *et al*. 2025). Here, we show that FAMA expression peaks during the night (Figure S1, Table S2). HOS1 transcripts peak at zt16 (Figure S1, Table S2) which might be a gating mechanism of Arabidopsis. *Hos1* mutant plants have reduced freezing tolerance (Ishitani *et al*. 1998) which could in turn indicate that by peak expression in the night the plant reacts to lower temperatures which are usually perceived during the night. Temperature, light and core clock expression change at dusk and dawn, leading to the expectation of two timepoints of diurnal transcript peaks in downstream targets. Previous analyses with microarrays identify a strong peak in the morning for all diurnal transcripts under the same photoperiod (Bläsing *et al*. 2005). The RNA-seq data acquired here presents its peak in diurnal transcripts at the beginning and during the night (Figure 1D). Both the growth conditions which differ in this experiment compared to Bläsing *et al*. (2005) in lighting by LEDs vs. fluorescent light, entrainment (22°C/18°C) vs. constant temperature (20°C), soil vs. plates, 50% vs. 60-70% relative humidity, age and the technical differences with RNA-seq no longer biasing against very low abundant and very high abundance transcripts may have contributed to the observed differences. The peak-at-night pattern is especially present in kinases with strong peaks in transcript abundance at zt14 (Figure 2C), but also in E3 ligases and TFs. Kinases expressed in mature leaves also show the most spatially resolved genes (26.6%) compared to TFs (22.48%) and lastly E3 ligases with only 17.35%, indicating importance of phosphorylation for nuanced regulation in Arabidopsis.

The temporal expression pattern of core clock components has been analyzed in detail (reviewed in Staiger *et al*. 2013), while the spatial expression pattern has gained less attention. We show here that the core clock components like CCA1, LHY, or TOC1 are highly temporal (Figure S2) with expected peak phases (Staiger *et al*. 2013), but ubiquitously expressed in all cell types (Figure S1). This underlines that every cell expresses the components to keep the time and matches previous analyses demonstrating cell type specific circadian rhythms (Endo *et al*. 2014; Qin *et al*. 2025). Clock TFs LHY, CCA1, PRR7 and PRR9 are significantly differential in the phloem (Figure S1, Table S4), which could be linked to a dominant vasculature clock identified previously (Endo *et al*. 2014). The phloem generally has the most differentially expressed genes (Table S4). This may reflect an only partially recognized role of the phloem as a signaling hub or it may point to technical challenges with spatial transcriptomics. Spatial transcriptomics data are often skewed towards annotating higher abundance transcripts as differential and dependent on high quality cell type annotation to reduce noise (Lähnemann *et al*. 2020). To avoid bias towards high abundance transcripts, we added the fraction of cells with expression as an alternative measure (Table S1), which considers only on/off states for each gene per cell. This measure also captures known enrichments of HOS1 and FAMA in the guard cells but shows a more ubiquitous expression of many spatially differential transcripts including core clock regulators (Figure S1).

The proportion of spatially expressed genes is generally lower compared to diurnal numbers (Figures 2B, 3B, 4B). This could be a result of general underrepresentation of low abundance transcripts in single cell data (Lähnemann *et al*. 2020) and the transcripts of the regulatory genes examined here being of significantly low abundance (Figure 1C) further reducing statistical power. Since many transcripts also oscillate on a cell type level (Endo *et al*. 2014; Qin *et al*. 2025), sampling time of the single cell samples additionally limits transcript detection and needs to be taken into consideration for bulk and single cell experiments. For the single cell data reanalyzed here (Lee *et al*. 2025), no time of sampling relative to the 16h photoperiod was provided. Given that CCA1 and PRR7 are expressed in many cells of all cell types, and PRR9 as well to a lesser degree, but ELF3 barely and ELF4 completely absent (Figure S1), this best matches the observed clock transcript state shortly after lights turned on at zt2 in our time-course (Figure S2). Although long day time-course analyses result in overall similar diurnal transcript numbers (Redmond *et al*. 2025), length of day and temperature influence clock entrainment resulting in observable expression differences (Oravec & Greenham 2022).

We show that overall, transcripts of proteins mediating the intermediate layer of regulation are of lower transcript abundance compared to all other genes (Figure 1C). Using bulk RNA-seq with sufficient read depth, the data identified diurnal rhythmicity in all intermediate regulatory sets for 52% (E3 ligases), 58% (kinases) and 59% (TFs) of expressed genes, which is slightly below the 62% of all expressed genes. Combining both diurnal and spatial datasets, we are able to resolve 66.1% of kinases (Figure 2B), 57.3% of E3 ligases (Figure 3B) and 63.2% of TFs (Figure 4B) in time, space or both. This dynamically expressed intermediate regulatory layer provides ample room to maintain tight regulation of general metabolism, development and stress response via transcription, phosphorylation and protein degradation. The time-course integrated with spatial data serves as an accessible resource (Table S1) to the community to identify rhythmic expression pattern in gene sets of choice.

## Data Availability

Raw reads are available under ARC (https://git.nfdi4plants.org/sanja.zenker/ath_timecourse) and NCBI SRA Bioproject PRJNA1333938 (made available upon publication). For easy access of the time-course data, we provide two shiny applications for local execution within the ARC: one to visualize tpms of individual Arabidopsis gene identifiers over time and another one to generate a heatmap for individual gene sets. Both applications with processed data and the exemplary gene sets used in this study can be downloaded from the ARC (https://git.nfdi4plants.org/sanja.zenker/ath_timecourse).

## Acknowledgements

We thank Prisca Viehöver for sequencing and the CeBiTec compute cluster for computational resources. This work was supported by the de.NBI Cloud within the German Network for Bioinformatics Infrastructure (de.NBI) and ELIXIR-DE (Forschungszentrum Jülich and W-de.NBI-001, W-de.NBI-004, W-de.NBI-008, W-de.NBI-010, W-de.NBI-013, W-de.NBI-014, W-de.NBI-016, W-de.NBI-022). SZ was supported by the DFG (INST 86/2288-1 to AB). Figure 2A, 3A and 4A were created with BioRender.com.

## Author contributions

**Sanja Zenker**: Data Curation; Formal Analysis (lead); Investigation; Visualization; Writing – original draft, review & editing. **Katharina Schiller**: Data Curation; Formal Analysis (supporting); Investigation; Visualization; Writing – original draft, review & editing. **Andrea Bräutigam**: Conceptualization; Funding Acquisition; Project Administration; Resources; Supervision; Writing – original draft, review & editing.

## Supplementary Information

**Figure S1 Heatmap of commonly known examples of kinases, E3 ligases and transcription factors.** Heatmap showing expression (z-score tpm) of all diurnal transcripts and fraction of cells with expression for each cell type. Stars indicate significantly differential expression in the cell type. Single cell data reanalyzed from (Lee *et al*. 2025).

**Figure S2 Expression patterns of core clock regulators.** Transcripts per million (tpm) of all three replicates at each Zeitgeber timepoint. Mean values per timepoint are connected by a line.

**Table S1 Transcripts per million over time and expressed cell ratio for all genes.** Tpm values for all Arabidopsis loci mapped with kallisto on primary transcripts. Timepoint is indicated by zt (Zeitgeber time; hours after lights turned on) and triplicates are denoted by R1/2/3. Expressed cell ratio was calculated from single cell data of 21d old Arabidopsis rosettes reanalyzed from Lee et al. (2025) A single-cell, spatial transcriptomic atlas of the Arabidopsis life cycle. Nature Plants. https://doi.org/10.1038/s41477-025-02072-z.

**Table S2 Analysis of rhythmic genes from JTKcycle.** Output from JTK_CYCLE analysis on expressed genes (tpm>=1 in all three replicates for at least one timepoint).

**Table S3 Enriched GO terms over time.** GO term enrichments on genes with peak at given phase (JTK_CYCLE output, Table S2) calculated with topGO and filtered for q-value<0.05 (Benjamini-Hochberg corrected).

**Table S4 Differentially expressed genes in cell types**. Cell type specific genes as defined by findAllMarkers output from Seurat, filtered for markers in maximally two cell types and p_val_adj<0.05. Seurat object of snRNA-seq data of 21d old rosettes from GSE226097 and cell type annotations from Supplementary Table 2 were retrieved from Lee et al. (2025) A single-cell, spatial transcriptomic atlas of the Arabidopsis life cycle. Nature Plants. https://doi.org/10.1038/s41477-025-02072-z.

