## SupplementalFigures for "Time- and space-resolved regulatory circuits in *Arabidopsis thaliana*"

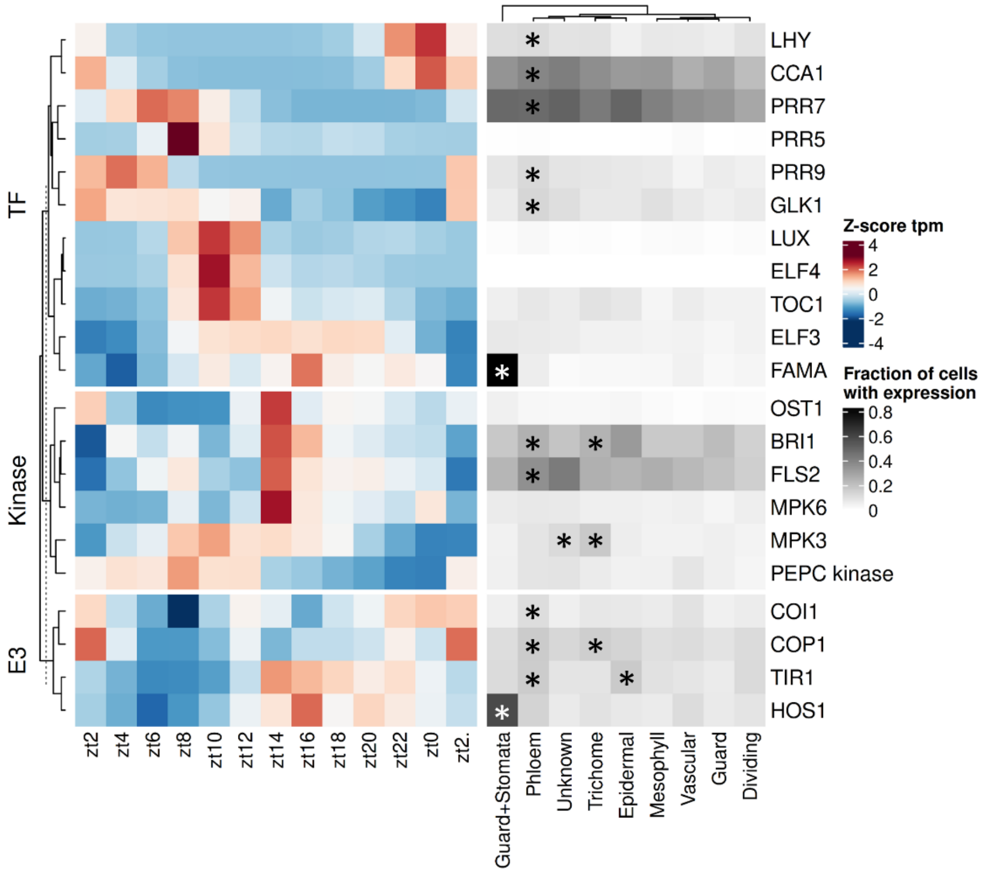


**Figure S1 Heatmap of commonly known examples of kinases, E3 ligases and transcription factors.** Heatmap showing expression (z-score tpm) of all diurnal transcripts and fraction of cells with expression for each cell type. Stars indicate significantly differential expression in the cell type. Single cell data reanalyzed from (Lee *et al.* 2025).

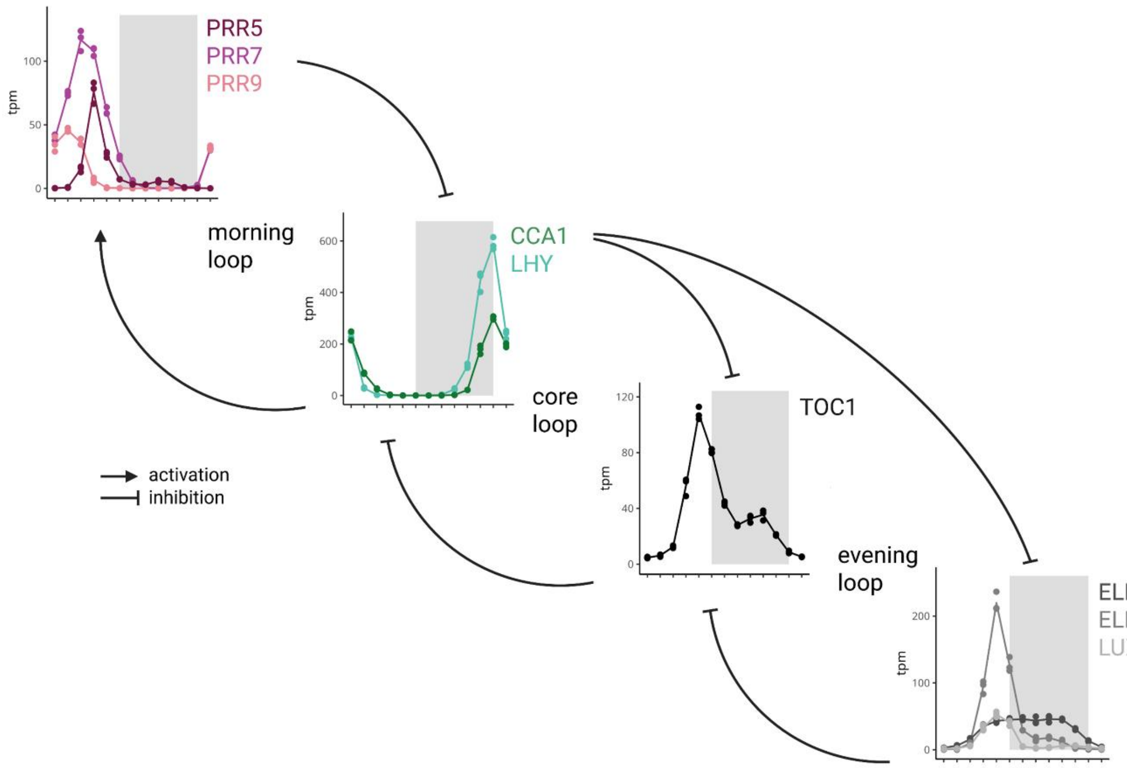


**Figure S2 Expression patterns of core clock regulators.** Transcripts per million (tpm) of all three replicates at each Zeitgeber timepoint. Mean values per timepoint are connected by a line.
